# β-Blocker inhibits myocardial infarction-induced brown adipose tissue D2 activation and maintains a low thyroid hormone state in rats

**DOI:** 10.1101/353052

**Authors:** Fernando A. C. Seara, Iracema G. Araujo, Güínever E. Império, Michelle P. Marassi, Alba C. M. Silva, André S. Mecawi, Luis C. Reis, Emerson L. Olivares

## Abstract

Considering the recognized role of thyroid hormones on the cardiovascular system during health and disease, we hypothesized that type 2 deiodinase (D2) activity, the main activation pathway of thyroxine (T4)-to-triiodothyronine (T3), could be an important site to modulate thyroid hormone status, which would then constitute a possible target for β-adrenergic blocking agents in a myocardial infarction (MI) model induced by left coronary occlusion in rats. Despite a sustained and dramatic fall in serum T4 concentrations (60-70%), the serum T3 concentration fell only transiently in the first week post-infarction (53%) and returned to control levels at 8 and 12 weeks after surgery compared to Sham group (P<0.05). Brown adipose tissue (BAT) D2 activity (fmoles T4/min.mg ptn) was dramatically increased by approximately 77% in the 8th week and approximately 100% in the 12th week in the MI group compared to that of the Sham group (P<0.05). Beta-blocker treatment (propranolol given in the drinking water, 0.5 g/L) maintained a low T3 state in MI animals, dampening both BAT D2 activity (44% reduction) and serum T3 (66% reduction in serum T3) compared to that of the non-treated MI group 12 weeks after surgery (P<0.05). Propranolol improved cardiac function (assessed by echocardiogram) in MI group compared to MI-non treated one by 40 and 57 % 1 and 12 weeks after treatment respectively (P<0.05). Our data suggest that the beta-adrenergic pathway may contribute to BAT D2 hyperactivity and T3 normalization after MI in rats. Propranolol treatment maintains low T3 state and improves cardiac function additionally.

## 1. Introduction

Ischemic heart disease remains the leading cause of death worldwide (1). Overall prognosis has been aggravated by limitations of available therapies and poor cardiac healing capacity. These limitations have fueled the search for alternative therapeutic strategies with focus in non-classical systems implicated in heart failure pathophysiology (HF), such as thyroid system (2).

One of these promising “non-classical” systems is the thyroid system. Thyroid hormones (TH) play important roles in cardiovascular homeostasis. Although many studies have previously described the cardiovascular effects of TH during health and disease (3,4), the physiological repercussions of myocardial infarction (MI) on thyroid status remains elusive so far. Surprisingly, the majority of studies have been performed in humans. Most of them have shown low serum triiodothyronine (T3) and no changes in thyrotropin (TSH) and thyroxine (T4) levels, a condition called “low T3 syndrome” (5,6). Although some studies have suggested that impaired T4-to-T3 conversion by deiodinases could explain the low T3 state elicited by MI (7,8), clinical evidences can be limited by the unfeasibility of performing a controlled evaluation over time due to different outcomes and drugs used. As a result, the mechanisms underlying possible changes in plasma TH concentration during HF development remain unclear.

Either systemic or cardiac hypothyroidism elicited by HF can be associated with the induction of ectopic cardiac type 3 deiodinase (D3) activity, the main inactivating pathway of TH (9,10). These features confirm systemic and local hypothyroidism induced by ectopic D3 activity, as well as the pathophysiology of “consumptive hypothyroidism,” which has been previously described in patients with large D3-expressing tumors (11), and do not support the “low T3-syndrome” model proposed in patients with HF (7,8). Even so, despite the persistent decrease of circulating T4 levels, T3 levels can return to basal levels in late phases of MI, i.e., 8 and 12 weeks post-surgery (12). Although high expression levels of ectopic cardiac D3 activity have been proposed to explain hypothyroidism following MI, the mechanism by which T3 can otherwise be normalized after MI remains unclear. Because type-1 deiodinase (D1) activity remains low throughout 12 weeks of MI, the type 2 deiodinase (D2) pathway has become the potential mechanism underlying, at least in part, the progressive re-establishment of serum T3 after MI (12).

Both D1 and D2 can convert T4 into T3 and, although D2 is crucial for local T3 production, it can also affect circulating T3 levels (13–16). In rats, either Dio2 mRNA expression or D2 activity have been mostly observed in the anterior pituitary, cerebral cortex (17), and brown adipose tissue (BAT) (18), though it has also been detected in thyroid and skin (19,20). Thus, unsurprisingly, BAT has been widely investigated as a potential source of circulating T3 (21–24). Together, these physiological properties have fueled studies to better understand the roles of BAT on thyroid homeostasis over cardiovascular diseases and to seek new pharmacological approaches for HF patients.

It has been demonstrated that D2 activity can be modulated by several factors, including sympathetic stimulation (25,26). Because MI and HF can induce sympathetic overactivation (27) and hypothyroidism (12), we hypothesized that D2 activity can be increased in response to MI. Although the role of sympathetic overactivity in the pathophysiological progression of HF, as well as the therapeutic benefits of beta-adrenergic blockers have been widely demonstrated, there have been little evidences on the repercussion of these drugs on BAT D2 activity. Therefore, we aimed to investigate whether i) the D2 pathway is overactivated following MI and ii) treatment with propranolol (the first successful beta-blocker developed) can influence the thyroid hormone economy and somehow partly explain its beneficial effects in a MI model.

## 2. Material and Methods

### 2.1 Animals

Male Wistar rats (200-250 g) were maintained in cages (4 per cage) in a room under controlled temperature (24 ± 2 °C) and lighting (lights on from 6:00 to 18:00 h), with free access to food and water. Animal handling and experimental procedures were performed according to the Guide for the Care and Use of Laboratory Animals published by the US National Institutes of Health (NIH Publication No.85-23, revised 1996) and the institutional committee of ethics and animal welfare (CEAW number: 23083.004836/20120-58).

### 2.2 Experimental Myocardial Infarction

MI was induced following the procedure previously described and modified by our group (JOHNS & OLSON, 1954; Olivares et al., 2007a). After anesthesia (Isoflurane - Biochimico^®^, Rio de Janeiro, Brazil), a skin incision was performed at the left parasternal level, followed by dissection of the pectoralis major and minor muscles. The incision was made between the 4th or 5th left intercostal spaces, through which the heart was externalized. Left coronary artery was located and connected as close as possible to its origin on the aorta. Hearts were then quickly placed in its original anatomical position. Sham-operated group (Sham) was subjected to the same surgical procedure as the infarcted group but without left anterior coronary artery occlusion. Prophylactic doses of veterinary antibiotic (0.2 mL, im, Pentabiótico Veterinário Pequeno Porte^®^, Fort Dodge, Brazil) and analgesic flunixin meglumine (2.5 mg/kg, im, Banamine^®^, Schering-Plough, Brazil) were given.

### 2.3 Heart Function Assessment and Pathology

Electrocardiogram (ECG) was registered one day after MI (Miranda et al., 2007) and echocardiogram (ECHO) evaluations were performed 1, 6 and 12 weeks after surgery to assess cardiac function. Echocardiograph color-system (Megas/Esaote) equipped with a 10 MHz electronic-phased-array transducer was used. Images were obtained from the left parasternal and apical windows. Short-axis 2-dimensional views of LV were taken at the level of papillary muscles to obtain the M-mode recordings. Systolic function was expressed by the ejection fraction (EF %), calculated by Simpson’s method, after the LV volume calculation: systolic and diastolic LV long axis were measured on long-axis view, and systolic and diastolic LV short axis, traced at the level of papillary muscle, were measured on transversal view.

Pathological analysis was performed as previously described (Olivares et al., 2004). Heart weight (HW), lung weight (LW) and liver weight (LiW) relative to body weight (BW) were calculated. The percentage of scar tissue in LV was calculated as described previously (Spadaro et al., 1980). Hearts were perfused with 4% paraformaldehyde in phosphate buffer. LV samples were sliced from apex to base (1–2 mm), and the slices were labeled as A (at the apex), B, C and D. Hematoxylin-eosin and Picrosirius staining were performed in representative sections obtained from slice C, described as the most representative of the total infarcted length using an Axiovert 100 microscope (Zeiss Inc. Germany). Sections stained with Picrosirius were recorded with a digital camera and stored for later analysis. All digital files were analyzed with ImageJ software (version 1.27 z, National Institutes of Health, USA). The percentage of the infarcted endocardium and epicardium was calculated, and the average percentage infarct size was estimated.

### 2.4 Radioimmunoassay (RIA) for T3 and T4

Serum T3 and T4 were determined by specific Coated-Tube RIA kits (T3: DLS – 3100 Active TX, EUA; T4: DLS – 3200 Active TX, EUA). All procedures were performed following the recommendations of the respective kit.

### 2.5 D2 Activity

D2 activity was determined using methods previously described (Fortunato et al. 2008). Pituitary gland or BAT (40 mg) were homogenized with 0.5 mL (pituitary glands) or 1 mL (BAT) of 0.1 M sodium phosphate buffer containing 1 mM EDTA, 0.25 M sucrose, and 10 mM dithiothreitol, pH 6.9. Each homogenate sample (100 μL containing 150 μg protein for BAT samples and 50 μg for pituitary samples) was incubated for 3 hours at 37 °C with 1 nM [125I] T4 (Perkin Elmer Life and Analytical Sciences, Boston, MA, USA), 20 mM dithiothreitol and 1 mM propylthiouracil (PTU) in 100 mM potassium phosphate buffer, pH 6.9, containing 1 mM EDTA.

Total reaction volume was 300 μL. The reaction was stopped at 4°C in ice-bath followed by addition of 200 μL fetal bovine serum (Cultilab, BR) and 100 μL trichloroacetic acid (50%, vol/vol), and vigorous agitation. The samples were centrifuged at 8000 g for 3 minutes, and an aliquot of supernatant (360 μL) was collected to measure 125I liberated during deiodination reaction. Protein concentration was measured by the Bradford method after incubation of homogenates with NaOH (2.5 N) (Bradford, 1976). Enzyme activity was expressed as femtomoles of T4 deiodinated per min per milligram of protein.

### 2.6 Experimental design

#### Phase I) Heart function, T3 and T4 serum concentrations and D2 activity in male rats subjected to experimental MI

Wistar rats underwent MI (MI group, n = 20) or sham operation (Sham group, n = 18) and serial blood sample collections (1, 8 and 12 weeks after surgeries) through the jugular vein to measure serum T3 and T4 concentrations. ECHO exams were performed on the first and twelfth week post-surgeries, immediately before blood collection. At the end of one, eight and 12 weeks after the surgical procedures, the animals were randomly euthanized by decapitation after sedation as previously described (n = 4-10/time/group) and the tissues (BAT and pituitary gland) were collected for type 2 deiodinase activity measurement.

#### Phase II) Influence of chronic β-adrenergic blockade on heart function, T3 and T4 serum concentrations and D2 activity in male rats subjected to experimental MI

Wistar rats underwent MI (n = 32) or Sham (n = 28) procedures and were randomly assigned to four groups: rats treated daily with propranolol hydrochloride (Sigma Aldrich), a β-adrenergic antagonist in drinking water (0.50 g/L): Sham + Prop (n = 14) and MI + Prop (n = 16) or vehicle: Sham + water (n = 14) and MI + water (n = 16), during 1 (n = 7 – 8/group) or 12 (n = 7 – 8/group) weeks after surgeries. At the end of each protocol, all groups underwent ECHO analyses and were euthanized the next day. Blood samples and BAT were collected for T4 and T3 serum measurement by radioimmunoassay and type 2 deiodinase activity assessments, respectively.

### 2.7 Statistical Analyses

Data are expressed as the mean ± SEM. Statistical analyses were performed by unpaired Student “t” test or one-way ANOVA followed by Bonferroni multiple comparison post-test, using GraphPad Prism^®^ 4 (GraphPad Software, Inc., San Diego, USA). Pearson’s product moment correlation was used to assess the relationship between the D2 activity and the T3/T4 ratio in each period, and a Student “t” test for paired data was used to assess the significant differences. Differences were considered statistically significant for p < 0.05.

## 3. Results

### 3.1 Phase I

#### 3.1.1 ECG and Heart Function Assessment

ECG and ECHO validated the efficiency of the MI model. As expected, MI was characterized by rightward deviation of frontal QRS axis (ÂQRS), Q wave was observed in L1 and QRS index (I-QRS) decreased (Figure 1A). Animals that did not show ECG signs of MI (two animals in this study) were excluded from the study. ECHO (Figure 1B) showed sustained reduction (approximately 40-50%) of LV EF in all observed periods in infarcted animals compared to Sham group.

**Figure 1:**
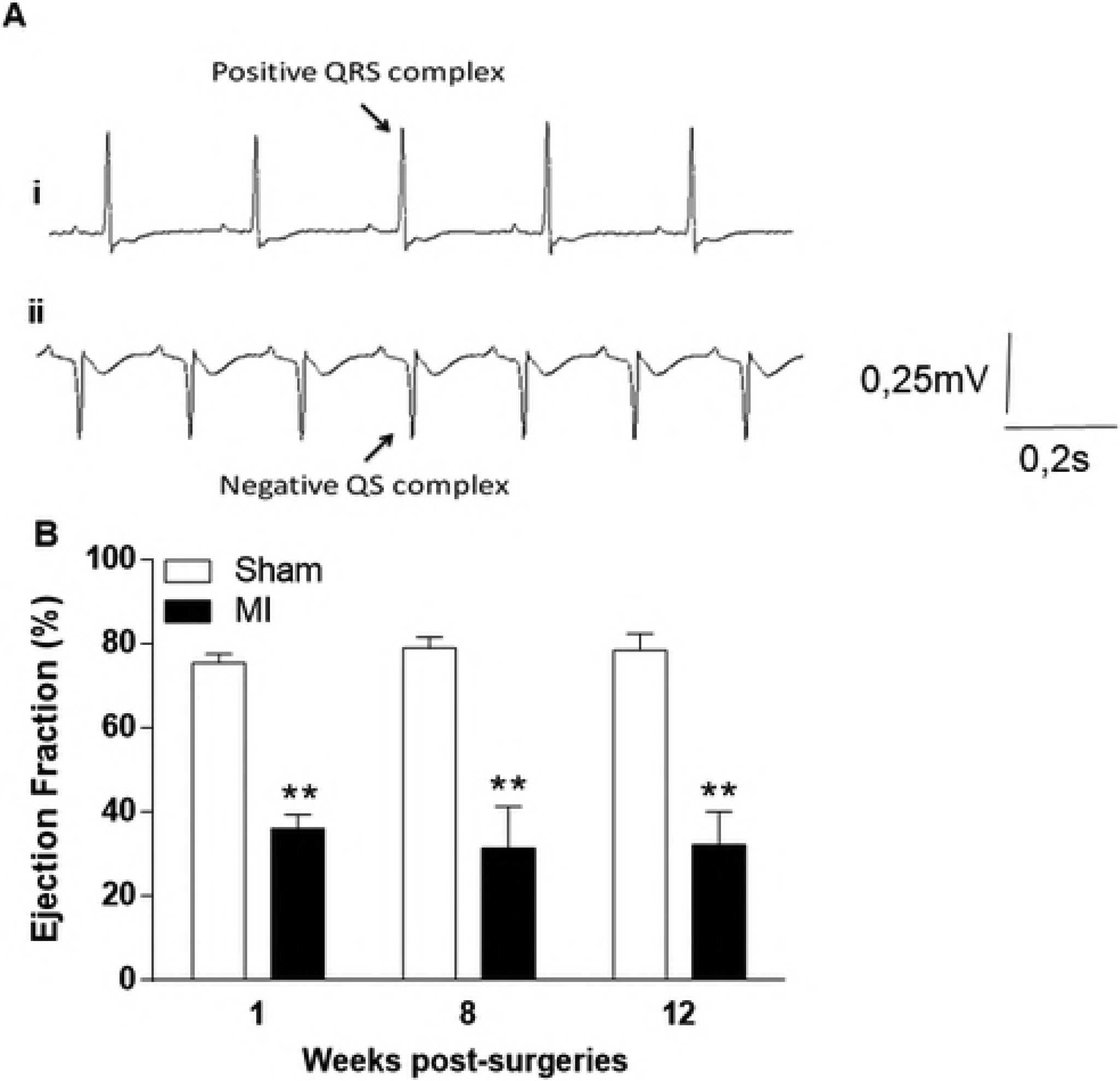
Characterization of MI-induced HF model. Representative ECG (A) from one Sham animal (i) and one MI animal (ii) and EF (%) progression of Infarcted and Sham-operated groups 1, 8 and 12 weeks after surgeries (B). Note the negative QRS complex and prominent Q-pathological wave (QS complex) exhibited by the infarcted animal (Aii). In B, the data are expressed as the mean ± SEM. **p < 0.01 vs. respective Sham group at the same week.

#### 3.1.2 Radioimmunoassay (RIA) for T3 and T4

Despite sustained decrease of serum T4 concentrations (approximately 60-70%) of MI compared to Sham group since the first week post-surgery (Figure 2A), serum T3 concentration fell only transiently in the first week (p < 0.01) and returned to control levels at 8 (p > 0.05) and 12 weeks (p > 0.05) after surgery (Figure 2B). Additionally, T3/T4 ratio was increased in MI rats versus Sham group at 8 and 12 weeks after surgery (Figure 2C). T3/T4 ratio was also higher in the 8th and 12th week compared to values observed in the MI group in the first week post-infarction (p < 0.01).

**Figure 2:**
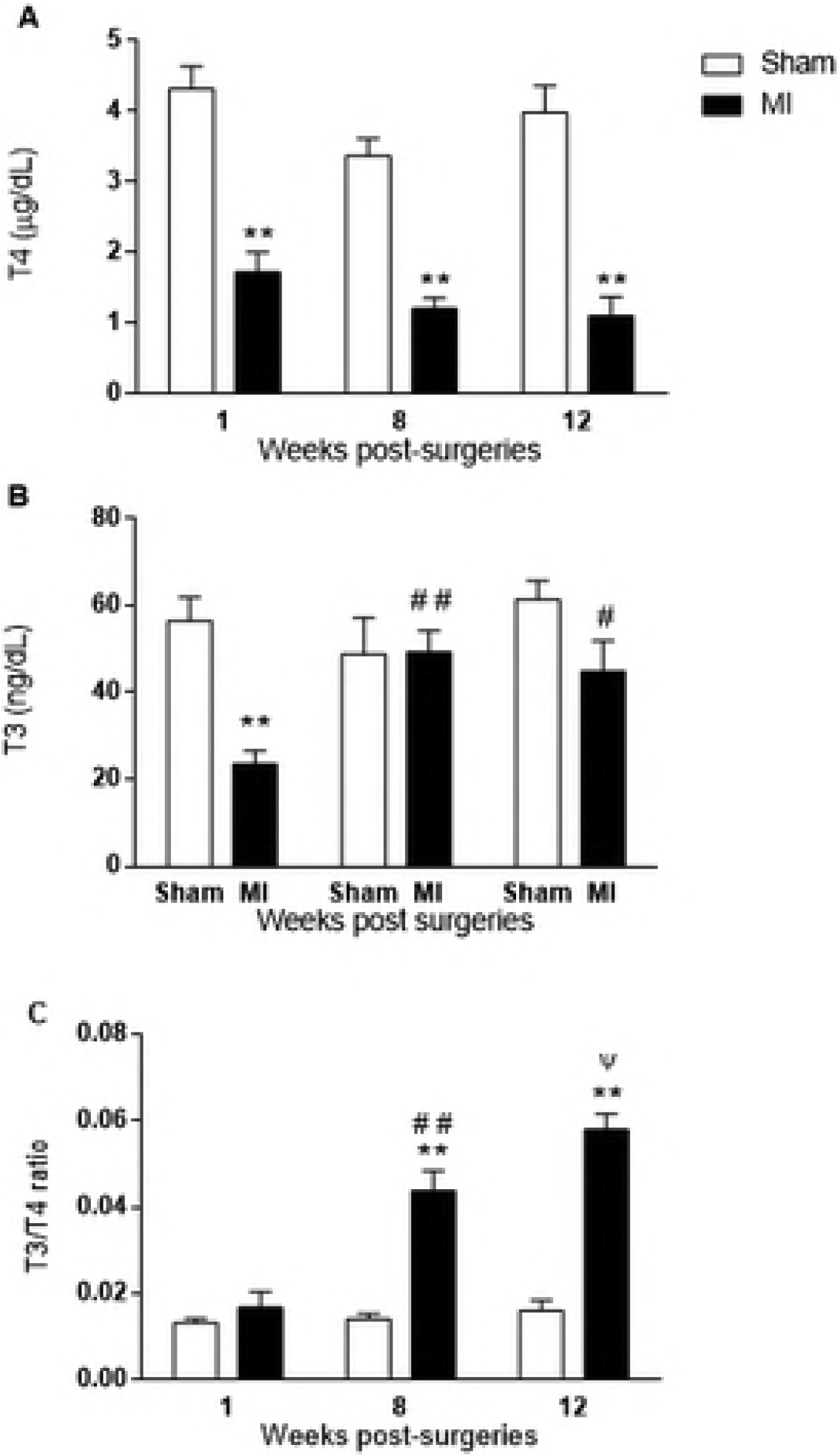
T4 (A) and T3 (B) serum concentrations and T3/T4 ratio (C) from Sham and infarcted (MI) groups 1, 8 and 12 weeks post-surgery. The data are expressed as the mean ± SEM. **p < 0.01, vs. respective control group at the same week; #p < 0.05 and ##p < 0.01 vs. MI group one week after surgeries; Yp < 0.05 vs. MI group 8 weeks after surgery.

#### 3.1.3 Type 2 Deiodinase Activity

BAT D2 activity (Figure 3a) increased 8 weeks (p < 0.01) and 12 weeks (p < 0.05) after MI versus Sham group, with no differences in the first week (p > 0.05). D2 activity progressively increased only in infarcted rats (Figure 3B), whereas no changes in pituitary D2 activity were noted (data not shown). Statistical correlation revealed strong and positive correlation between the T3/T4 ratio and BAT D2 activity in the MI group (p < 0.0001) over the course of the 12 weeks after the permanent coronary occlusion (Figure 3B).

**Figure 3:**
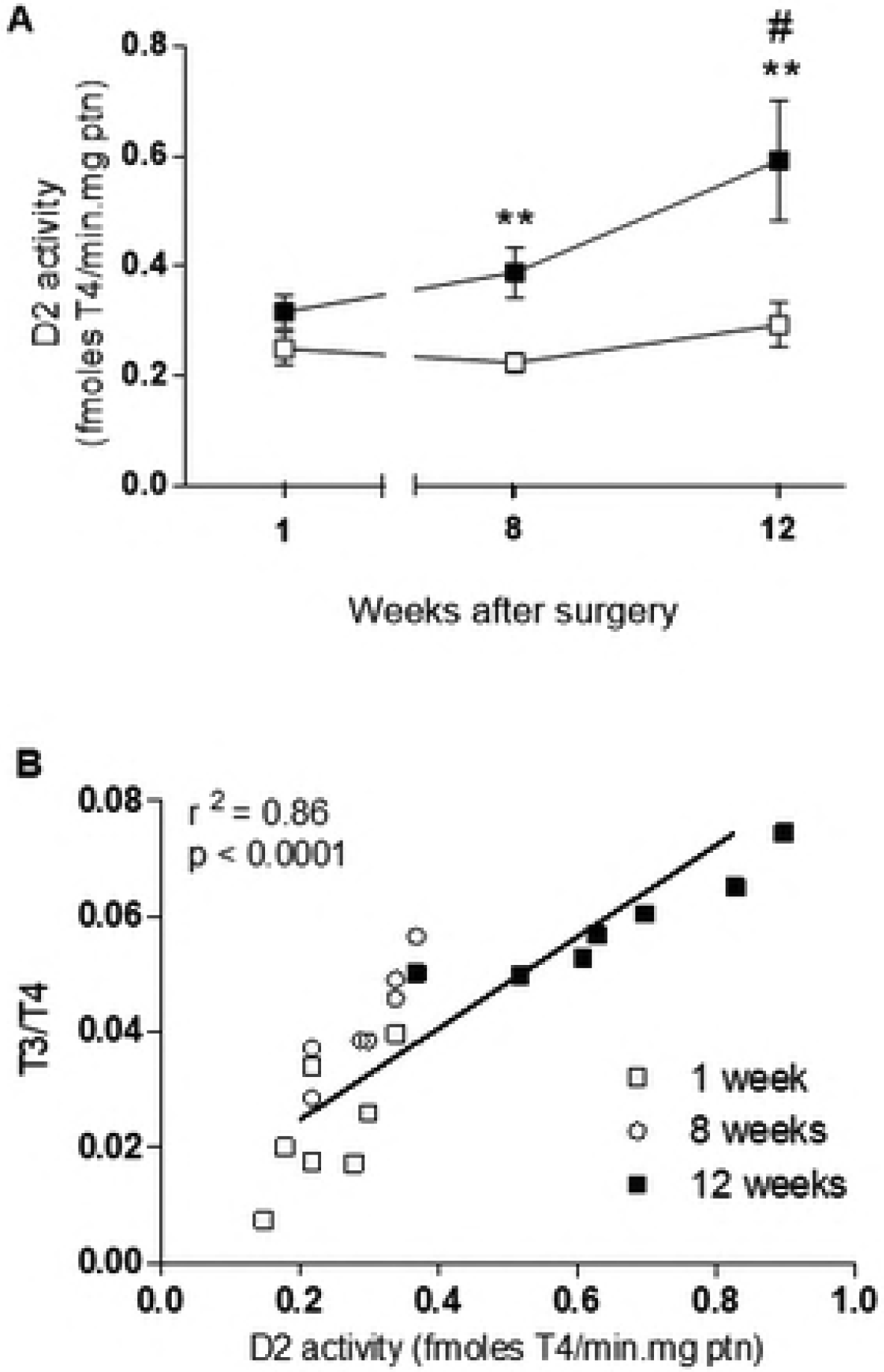
Temporal effects of MI on D2 activity. Progression of D2 activity in BAT (A) and pituitary gland (B) and correlation between the T3/T4 ratio and D2 activity in BAT 1 (white square), 8 (white circle) and 12 (black square) weeks from sham-operated (Sham) or infarcted rats over the 12 weeks of the experimental protocol. Each symbol in Figure C corresponds to the point of intersection between the D2activity in BAT and the T3/T4 ratio for each animal studied. r^2^ = Pearson correlation coefficient. The data are expressed as the mean ± SEM. **p<0.01 vs. respective control group; # p<0.05 vs. MI group in the first week.

### 3.2 Phase II

#### 3.2.1 Heart Function Assessment and Pathology

Propranolol prevented MI-induced tachycardia at 8 and 12 weeks post-surgery (P < 0.05), suggesting that cardiac adrenergic stimulation was blunted. ECHO analysis (Figure 4) demonstrated sustained reduction of LV EF (approximately 40-50% decrease) in both periods of observation in MI compared to Sham group. Cardiac function was improved by propranolol, including LV EF one and 12 weeks post-MI (Figure 4). However, this improvement was greater at 12 wks (57%) than 1 wk (40%) post-MI.

**Figure 4:**
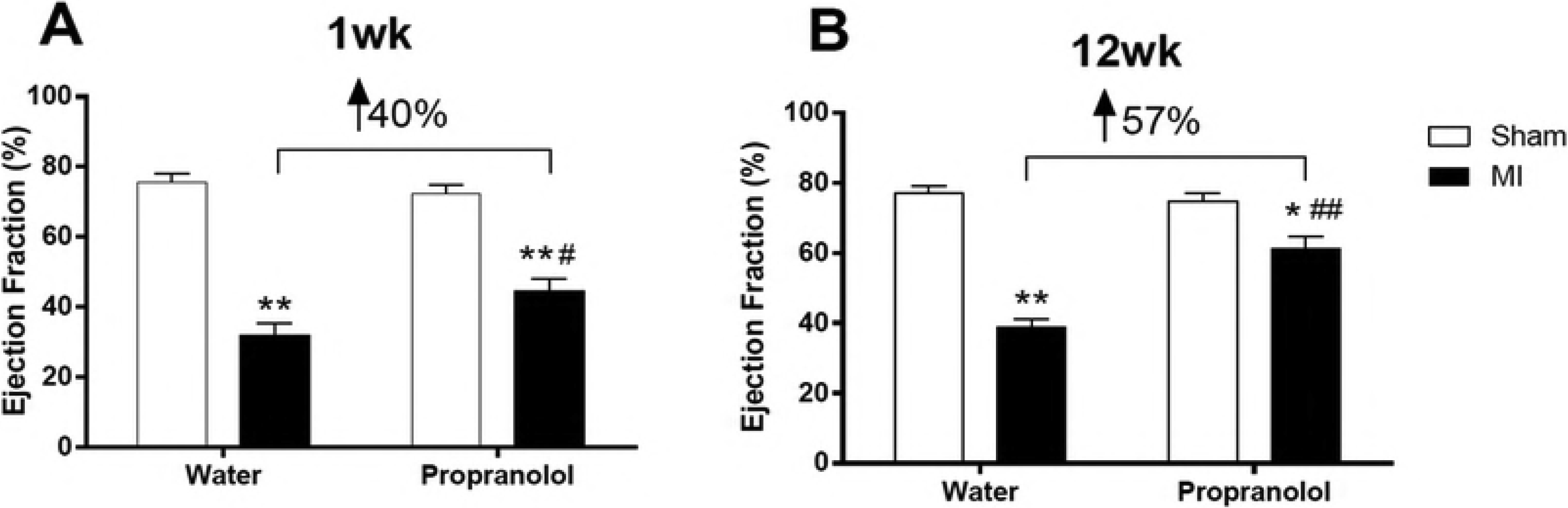
Effect of propranolol treatment on heart function assessed by echocardiography. LV EF (%) at 1 wk (A) and 12 wks (B) post-MI were compared between vehicle or the propranolol-treated sham group (white boxes) and MI group (black boxes). The data are expressed as the mean ± SEM. *p < 0.05 and **p < 0.01 vs. respective control group at the same week. #p<0.05 and #p<0.01 vs. MI + water group.

Pathology data (Table 1) showed that HW was significantly increased in the MI+Water group compared to the Sham+Water group at 1 and 12 weeks after infarction surgery (P < 0.05), suggesting the occurrence of cardiac hypertrophy. Conversely, the MI+Propranolol exhibited no increase of HW in the first week and a slight increase smaller than that observed in the first week compared to control (P < 0.05). Increased LW was also evident in MI+Water at both time periods, consistent with congestive failure, which was less apparent in MI Propranolol-treated animals. Again, propranolol treatment was more effective in the first week compared to 12 weeks post-MI. LiW did not change throughout the experimental protocol in all groups.

**Table 1.** Body weight and heart, lung and liver relative weight and infarct size at 1 and 12 wk post-surgeries.

| Groups | Parameters at Week 1 |  |  |  |  |
| --- | --- | --- | --- | --- | --- |
|  | Body weight (g) | Relative heart weight (mg/g) | Relative lung weight (mg/g) | Relative liver weight (mg/g) | Infarct size (%) |
| Sham + water | 260.9 ± 3.18<sup>a</sup> | 3.4 ± 0.14<sup>a</sup> | 5.2 ± 0.18<sup>a</sup> | 35.9 ± 1.06<sup>a</sup> | - |
| MI + water | 264.4 ± 3.33<sup>a</sup> | 5.7 ± 0.23<sup>b</sup> | 7.1 ± 0.17<sup>b</sup> | 37.7 ± 1.32<sup>a</sup> | 44.38 ± 1.47<sup>a</sup> |
| Sham + propranolol | 261.6 ± 2.73<sup>a</sup> | 3.6 ± 0.15<sup>a</sup> | 5.4 ± 0.14<sup>a</sup> | 35.6 ± 1.59<sup>a</sup> | - |
| MI + propranolol | 266.4 ± 3.53<sup>a</sup> | 4.3 ± 0.15<sup>c</sup> | 6.1 ± 0.10<sup>c</sup> | 39.4 ± 1.62<sup>a</sup> | 35.88 ± 1.82<sup>b</sup> |
| Groups | Parameters at Week 12 |  |  |  |  |
|  | Body weight (g) | Relative heart weight (mg/g) | Relative lung weight (mg/g) | Relative liver weight (mg/g) | Infarct size (%) |
| Sham + water | 450.3 ± 3.91<sup>a</sup> | 4.0 ± 0.09<sup>a</sup> | 5.8 ± 0.16<sup>a</sup> | 37.4 ± 1.16<sup>a</sup> | - |
| MI + water | 467.3 ± 3.87<sup>a</sup> | 6.2 ± 0.30<sup>b</sup> | 8.4 ± 0.13<sup>b</sup> | 39.3 ± 1.30<sup>a</sup> | 50.86 ± 2.20<sup>a</sup> |
| Sham + propranolol | 454.4 ± 4.36<sup>a</sup> | 4.2 ± 0.21<sup>a</sup> | 5.5 ± 0.16<sup>a</sup> | 38.2 ± 1.37<sup>a</sup> | - |
| MI + propranolol | 461.4 ± 3.35<sup>a</sup> | 5.5 ± 0.15<sup>b</sup> | 7.4 ± 0.20<sup>c</sup> | 41.7 ± 1.52<sup>a</sup> | 48.38 ± 1.82<sup>a</sup> |

Different letters indicate values significantly different ( $P < 0.05$ ).

Propranolol reduced the infarct size at the first week post-MI (P < 0.05) but not 12 wks post-MI (P > 0.05) as observed in MI+Propranolol vs. MI+Water at both times. These combined data suggest that cardiac hypertrophy, congestive HF and infarct size do not explain the improvement in cardiac function observed in the myocardial infarcted animals treated with propranolol at 12 wks. Therefore, other factors should explain the improvement in cardiac function observed in propranolol-treated infarcted rats, particularly in the late phase of cardiac failure.

#### 3.2.2 Radioimmunoassay (RIA) for T3

Serum T3 levels observed in MI-animals in this Phase (Figure 5) reproduced the results shown in Phase I (Figure 2). T3 concentrations fell transiently in the first week post-infarction (approximately 67% decrease) and returned to control levels at 12 weeks. Propranolol did not alter the decrease in T3 levels at 1 wk post-MI (approximately 57% decrease), but it was otherwise effective preventing plasma T3 restoration observed in MI-treated rats 12 weeks post-MI (Figure 5). The MI group treated with the β-blocker had values resembling hypothyroid status as observed in non-treated MI group at 1-week post-MI (approximately 66% decrease in T3 levels). Altogether, these data suggest that propranolol treatment for 12 weeks prevented the reestablishment of euthyroid status normally observed in response to MI.

**Figure 5:**
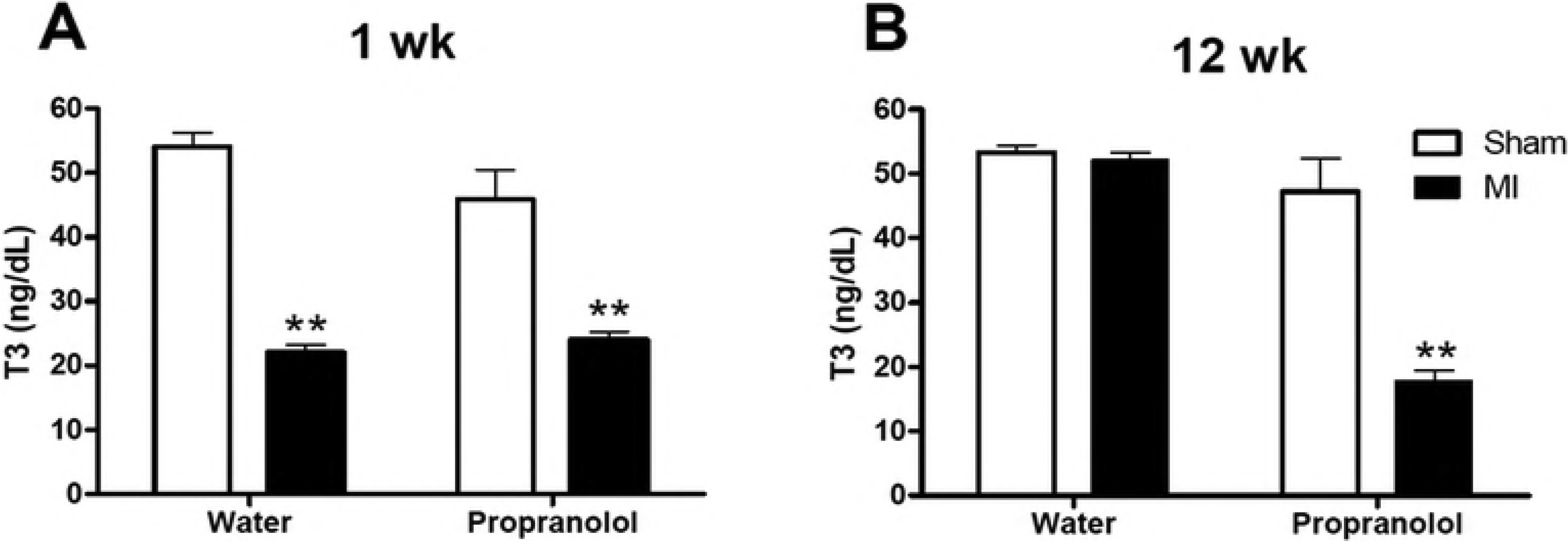
Circulating T3 levels at 1 wk (A) and 12 wks (B) post-MI were compared between vehicle or the propranolol-treated sham group (white boxes) and MI group (black boxes). The data are expressed as the mean ± SEM. **p < 0.01, vs. respective control group at the same week.

#### 3.1.3 Type 2 Deiodinase Activity

As expected, BAT D2 activity increased in MI+Water compared to Sham+Water group at 12 weeks (Figure 6B, p < 0.01). MI+Propranolol showed no difference compared to Sham+Propranolol at 12 weeks after the permanent coronary occlusion (Figure 6B, p > 0.05). No differences were observed among the groups at 1-week post-surgeries (Figure 6A, p > 0.05).

**Figure 6:**
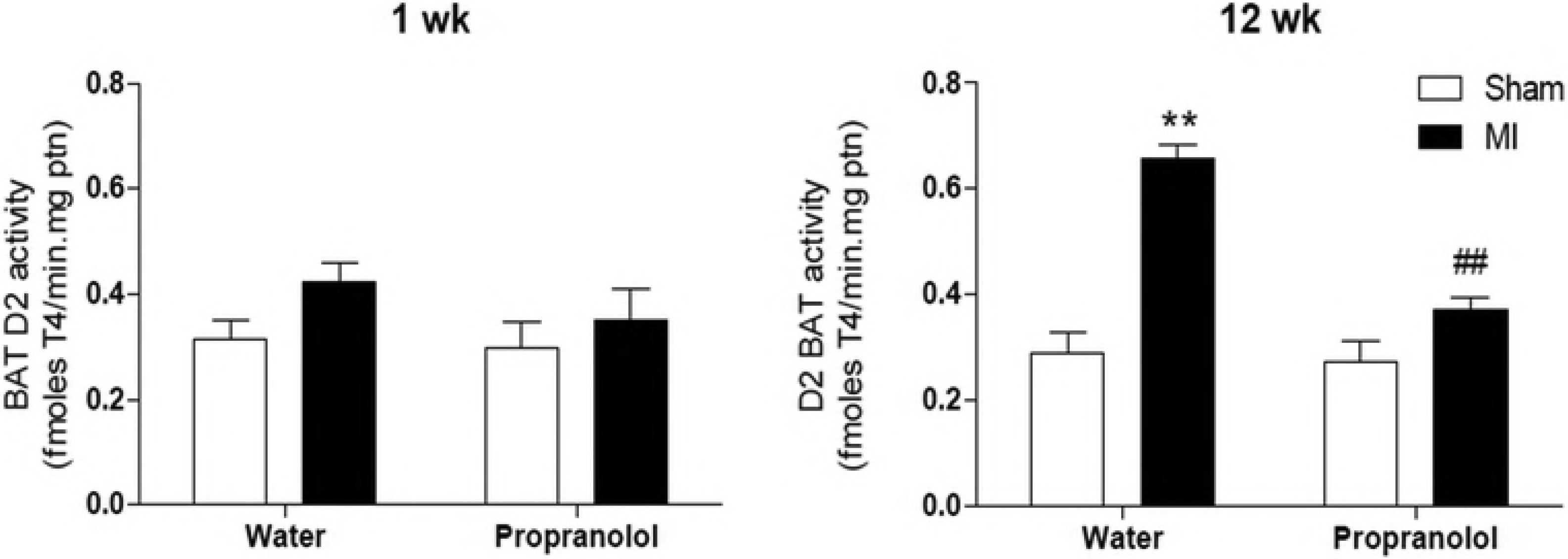
BAT type 2-deiodinase activity 1 (A) and 12 (B) weeks from sham-operated and infarcted rats treated or not with propranolol. **p<0.01 vs. respective control group; ## p<0.01 vs. MI group in the first week.

## 4. Discussion

In this study, we reported a progressive enhancement of D2 activity in BAT from rats after MI. The increased T4 to T3 conversion could correspond to an important mechanism responsible for serum T3 normalization observed at the late phase of MI, for previous evidences have suggested that D2 might also be important for serum T3 supply, besides its crucial role in intracellular T3 provisions (13,17,28–30).

D2 activity can be modulated by several factors, such as hypothyroidism, α- and β- adrenergic activation, and bile acids (14,25). Interestingly, propranolol did not prevent HF-induced hypothyroidism, but it prevented the normalization of serum T3 levels at the late phase of HF. As expected, these data suggest that β-adrenergic pathway plays a crucial role on BAT D2 hyperactivity and serum T3 normalization after MI. In such a condition, sustained decrease of serum T4 concentrations over 12 weeks after MI could lead to higher BAT D2 activity. In normal conditions, 50% of the intracellular T3 comes from deiodination processes induced by D2 (31), and a drastic reduction of circulating T4 levels would decrease intracellular T3 levels, consequently elevating BAT D2 activity (28). Despite reductions of circulating T3 and T4 levels (50% and 90% respectively), normal cardiac T3 levels have been demonstrated in model of iodine deficiency in rats (32). Accordingly, local conversion of TH can be increased by low circulating T4 levels, and BAT can increase as an alternative source for circulating T3 (33,34).

Because MI can be followed by autonomic imbalance, increased BAT D2 activity could be secondary to sympathetic nervous system overactivity (27). Both α and β-adrenergic receptors can be expressed by brown adipocytes (35), and the enhanced sympathetic activity can effectively modulate BAT metabolism, particularly D2 activity, as observed in cold-stress exposure (36,37). Because sympathetic overactivity can persist many weeks after permanent coronary occlusion (27), we suggest that in addition to hypothyroidism stimuli, increased BAT D2 activity could also be induced by the higher sympathetic activity towards BAT. Future studies employing surgical or pharmacological sympathectomy will be needed to further address this hypothesis.

Another potential mechanism to explain serum T3 normalization lies in D3 activity. If high cardiac D3 activity is associated with low serum thyroid hormones in the early phase of HF (38), T3 normalization observed in the late phase could be secondary to D3 inactivation. Additionally, we characterized and confirmed the consumptive hypothyroidism status, which has been previously described in patients with large D3-expressing tumors (11), as thyroxine injected early after MI could restore euthyroid status only 6 weeks after the beginning of the hormonal treatment. Indeed, there has been no evidence of persistent D3 induction in models of MI in rats (2). Future studies assessing a time course of D3 activity in this model are needed.

Interestingly, BAT D2 activity progressively increased after MI (Figure 3A), and it was positively correlated with the T3/T4 ratio (Figure 3C). The progressive increase in T4 to T3 conversion was expected, inasmuch as increased D2 activity can be associated with the longer half-life of D2 protein and increased Dio2 gene transcription (29,39). D1 activity could be affected by propranolol, since T4 to T3 conversion is also attributed to D1 activity. However, persistent low D1-activity 4 and 12 weeks after MI has already been described (12). Therefore, the possible effect of propranolol on D1-activity is not reasonable for D1 activity is already down-regulated in our model. Furthermore, it is also not rational to postulate a propranolol effect (block) on D3 activity, since D3 is a the main physiological inactivator of TH (40). At the same time, as far as we know, there is no description of beta-blockers on D3 activity in literature so far.

Since T3 was restored to control levels, though normalization of T3 was not associated with any improvement in cardiac function in infarcted rats, one may suggest that infarcted hearts were unresponsive to TH actions, consistent with previous evidences (41). Any differences in pituitary D2 activity was noted in the eighth and twelfth weeks after MI, despite the hypothyroxinemic status of infarcted rats (data not shown). Pituitary D2 can be affected by variations of T4 serum concentrations (28), which may be important for maintenance of D2 homeostasis (mainly in the cerebral cortex) and, thereby, intracellular T3 concentrations (30). Additionally, TSH secretion can be finely modulated by intra-pituitary D2-induced T4 to T3 conversion and by serum T3 levels (28,40). Some reports have also shown that large bolus of T4 can only inhibit TSH secretion if pituitary D2 activity is fully active, strongly supporting the concept of a crucial role of D2 in the control of TSH secretion in normal rats (42,43). Thus, in our model, normalization of serum TSH concentrations might be due to serum T3 normalization rather than intracellular conversion from T4, as previously described (12). In addition, the roles of other Dio2-positive tissues, such as skeletal muscle and heart on T3 restoration after MI remain elusive (44).

Beta-adrenergic blockers can improve survival in patients with HF, but their beneficial effects on TH economy during HF are elusive (45). Interestingly, early stage HF-induced hypothyroidism was not significantly affected by propranolol, though serum T3 normalization in the late stage of cardiac disease was prevented. Surprisingly, the most striking improvement of cardiac function observed in the MI group treated with propranolol occurred in the late stage (12 wks post-MI), and not the early stage, of HF (1 wk post-MI). Furthermore, cardiac hemodynamic improvement was not correlated with improvements of cardiac remodeling (infarct size and cardiac hypertrophy) or hemodynamic state (indirectly assessed by relative lung weight). Since MI was associated with hypothyroidism only in propranolol-treated rats at the end of 12 weeks, these data suggest that beta-blockers maintain a hypothyroid state, guaranteeing additional improvement in cardiac function.

The relationship between hypothyroidism and cardiac function has been previously described. Ueta et al. showed that a 10-day sympathetic overdrive worsened the phenotype of restrictive cardiomyopathy in D3KO mice (HtzD3KO) and lends itself as a model of cardiac-specific Dio3 inactivation and local hyperthyroidism (46). Consequently, LV dysfunction was further impaired, particularly diastolic properties, resulting in congestive HF and a higher mortality rate. In sharp contrast, wild type siblings could recruit myocardial D3, which resulted in adaptive cardiac remodeling and better cardiac function compared to D3KO mice. Furthermore, we recently demonstrated that hyperthyroidism furthered LV stunning while hypothyroidism was mostly followed by improved post-ischemic recovery of LV hemodynamic properties and decreased infarct size (47). These findings are consistent with our present work, which suggests that cardiac hypothyroidism is an adaptive condition important for guaranteeing an allostatic condition in HF.

## 5. Conclusion

In conclusion, our results suggest that the normalization of serum T3 concentrations after MI in rats can be induced by a progressive increase of D2 activity in BAT, which was potentially induced by the beta-adrenergic pathway post MI in rats. When this pathway was blocked by propranolol, the low thyroid hormone state persisted, and cardiac function was improved additionally, suggesting a new beneficial effect of beta-blockers in a HF model.

## Acknowledgements

None of the authors has any conflicts of interests. The authors are very grateful to Mr. Advaldo Nunes Bezerra, Mr. Ipojucan P Souza and Mr. Wagner Nunes Bezerra for the animal care and technical assistance. This research was supported by Conselho Nacional de Desenvolvimento Científico e Tecnólogico (CNPq) and Fundação de Amparo à Pesquisa do Estado do Rio de Janeiro (FAPERJ).

